# Reprogramming transcription factor networks with a HOX inhibitor to address Huntington’s disease biology beyond mutant huntingtin

**DOI:** 10.64898/2026.11.09.765359

**Authors:** Michael Klukinov, Rick Orr, William K. Martin, Shashidar Kori, David C. Yeomans, Julien Mamet

**Author notes:** Correspondence: David C. Yeomans; Julien Mamet.

## Abstract

Huntington’s disease (HD) is an inherited neurodegenerative disorder caused by CAG-repeat expansion in the huntingtin (*HTT*) gene. Although inherited CAG-repeat length and the underlying somatic instability are key determinants of age at onset, they explain only part of onset variability, suggesting additional disease mechanisms. We applied the Core Coefficient (CC), a transcription-factor (TF) network-decoding technology that quantifies TF functional dominance, to >100 post-mortem neuronal transcriptomes from HD and control brain datasets. Among human TF families, the HOX family is the only family that uniformly exhibits quiescence in controls versus large and significant functional dominance in vulnerable neurons of each HD donor brain we investigated. HOX TFs are already dominant at Vonsattel grade 0 and at inherited CAG-repeat length. The magnitude of HOX CC values correlates with age at onset and increases with disease grade and CAG-Age Product. HOX TFs form a self-sustained pathological unit targeting >4,000 genes implicated in neuronal patterning, identity, stress, signaling, structure, and senescence. In zQ175 mice, abrogation of mutant huntingtin (mHTT) protein production only transiently attenuates HOX dominance and does not durably eliminate it. COR-H800, a HOX inhibitor, restored TF-network hierarchy toward wild type, normalizing the functional dominance of transcription factors involved in neuronal differentiation, cell-fate determination, and developmental patterning, and improved disease-relevant phenotypes in R6/2 mice. These findings support the direct selective inhibition of HOX TFs to address the aberrant transcriptional state that can persist despite mHTT lowering or somatic instability-targeted therapeutic interventions.

## Introduction

Huntington’s disease (HD) is an inherited neurodegenerative disorder caused by CAG-repeat expansion in exon 1 of the huntingtin (*HTT)* gene, resulting in expression of mutant huntingtin protein (mHTT). While the inherited CAG tract length initially dictates disease risk, the repeat exhibits somatic instability, with progressive lengthening over an individual’s lifespan through a process termed somatic expansion.^1–3^ Despite the ubiquitous presence of the mutation throughout the body, this expansion preferentially affects striatal medium spiny neurons (MSNs) and subsets of cortical pyramidal neurons.^4,5^ Hence, mHTT exposure alone does not explain why particular neuronal populations enter dysfunctional and degenerative states and other cells do not. In various models of HD, altered corticostriatal synaptic circuit function can precede frank neuronal loss, indicating that neurons progress from dysfunction to terminal degeneration.^6–8^ Human HD tissue likewise shows widespread, cell-type-specific dysregulation of gene programs related to neuronal identity, synaptic maintenance, stress response, metabolism and inflammation.^9–12^

Altogether these data support the hypothesis that long-term transcriptional alterations fuel neuronal impairment and ultimately degeneration in HD. While dysregulation of approximately a dozen individual TFs has been associated with neurodegeneration in HD (e.g., REST/NRSF, p53, NF-κB, C/EBPα, HSF1, SREBF2 or ELK1)^13–19,^ three fundamental questions regarding how transcriptional dysregulation underlies the disease are unanswered:

- First, is there a TF or group of TFs that acts as a common denominator of HD pathogenesis across vulnerable neurons, disease grades and patients?
- Second, because transcriptional dysregulation is prone to memory and can acquire self-sustaining properties that persist after an initiating trigger has subsided, as observed in a number of pathological states (e.g., metabolic diseases, inflammation, chronic pain)^20–24^, to what extent does that common denominator remain dependent on mHTT abundance and CAG-repeat expansion over time?
- Third, can the blockade of this common denominator reverse transcriptional dysregulation and produce therapeutic benefits? To answer these questions, we used a systems biology approach that decodes TF network hierarchy and quantifies TF functional dominance, a property that is not measurable through differential gene expression analyses alone.

Using this approach, we demonstrated that the functional leverage of TFs over a cell’s function or state is poorly correlated with their expression level.^25^ This means that the general interpretation that an increase in expression implies a gain of functional activity of a gene, and vice versa, is limited for TFs. For instance, STAT5 is known to promote cell proliferation, but its overexpression does not alter proliferation.^26^ A low expression level can also mean increased dependence on a TF; for example, SPI1 is reduced in acute myeloid leukemia cells compared to normal hematopoietic cells, but leukemic cells depend on the residual expression of SPI1 for survival.^27^

An explanation for this uncoupling lies in the fact that TFs operate as functional networks with dominant and redundant components.^25^ These networks consist of individual TFs interacting with one another to collectively organize and regulate transcriptional programs.

We developed the Core Coefficient (CC) algorithm to quantify the functional importance of each TF within the network of a given cell type, with the goal of identifying functionally dominant regulators rather than simply the most abundant transcripts. We previously demonstrated the correlation between TF CC values and TF dominance in mechanistically distinct diseases, including cancer, fibrosis, and myocardial infarction, and experimentally validated this relationship.^25^

To characterize TF dominance in the context of HD, we built TF networks from post-mortem transcriptomes of vulnerable neurons and analyzed them agnostically with the CC. Results show a robust and systematic gain of functional dominance of a single family of TFs in HD that is normally dormant in the adult forebrain: the HOX family (HOXs). CC analysis of the networks shows that ectopic HOX activity dysregulates thousands of genes in MSNs associated with neuron dysregulation and degeneration. HOX dominance is already present at inherited CAG repeat length in HD and can uncouple itself from mHTT levels, offering new insights into the pathogenesis of the disease.

To demonstrate experimentally the potential of targeting HOXs to treat HD, we designed a decoy compound named COR-H800 that blocks HOX activity by mimicking the DNA motifs bound by HOX members on the genome. COR-H800 treatment eliminated HOX dominance in MSN networks and improved the phenotype of the transgenic R6/2 mouse model of HD. This work identifies the HOX family of TFs as a promising therapeutic target for HD, supporting a new direction beyond strategies focused on restraining somatic instability and lowering the abundance of mHTT.

## Results

### HOXs become functionally dominant in the TF networks of vulnerable neurons

To decode the functional hierarchy of TF networks in HD brains and identify a potential common denominator of transcriptional dysregulation, we first built MSN TF networks from a dataset that contains transcriptomic data from the putamen and caudate nucleus of HD and control donors (GSE152058)^28^ and analyzed them with the CC. Comparison of the functional network hierarchy revealed a striking phenomenon: the significant gain of functional dominance of the HOX family of TFs in HD (Figure 1A). Expanding the analysis to all 100 human TF families revealed a highly targeted network response wherein only the HOX family systematically gains functional dominance in HD (Figure 1A). This phenomenon is observed consistently in each patient analyzed in the dataset (Figure 1B). To further investigate the breadth of this HOX response, we built separate networks of D1 and D2 MSN subtypes for the two striatal regions. CC analysis of those networks confirmed a HOX response in D1 and D2 MSN subtypes with respective fold gains of ≈ 13 and ≈ 5 in the caudate region, and ≈ 36 and ≈ 3 in the putamen. To further confirm the commonality of the HOX response, we next built MSN networks using the nemo:dat-ztfn3cc dataset, which includes snRNAseq data from the caudate of 50 individuals with HD and 53 controls.^29^ The cohort was balanced by sex (29 control males, 24 control females, 25 HD males, 25 HD females) and spanned a broad range of ages at motor or cognitive onset (13 to 75 years), ages at donation (26 to >89 years) and disease grades (0 to 4). Results showed a HOX functional response in the MSNs of all persons with HD compared to controls (Figure 1C), confirming the commonality of HOX dominance.

**Fig. 1.**
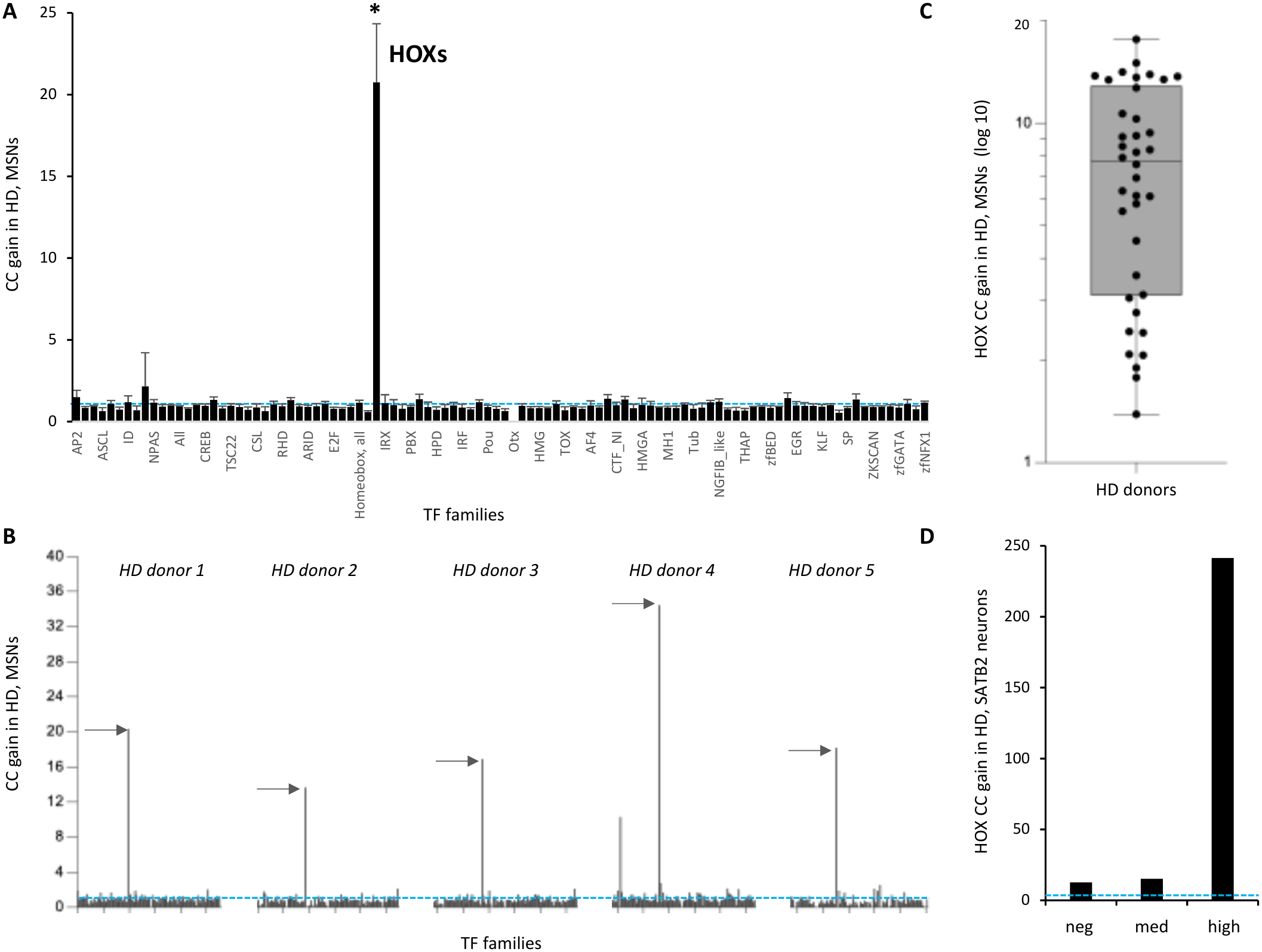
HOX TFs gain functional dominance in vulnerable neurons. (A) CC analysis of the 100 human TF families in networks constructed from postmortem transcriptomic data shows that the HOX family exhibits a significant increase in CC, reflecting increased functional dominance, in MSNs from the caudate and putamen. Data are presented as mean ± SEM (GSE152058 dataset; N = 8 controls and 5 HD donors). (B) Individual-donor analysis from the GSE152058 dataset. The HOX family is identified by a horizontal arrow. (C) HOX CC gain in MSNs from the caudate nucleus of individual HD donors in the nemo dataset with sufficient sequenced cells for TF-network construction (N = 53 controls and 38 HD donors). (D) HOX CC gain in SATB2-negative, SATB2-medium, and SATB2-high pyramidal neurons of layer 5a of the motor cortex (GSE233387 dataset; N = 6 controls and 13 HD donors). For control and HD donors, data were pooled to obtain sufficient cells for TF-network construction for each neuronal population; therefore, statistical comparisons were not performed for this panel. For all applicable panels, HD CC gain was calculated by normalizing the CC of each TF family in HD to the corresponding mean CC in controls from the same dataset. *p < 0.05, Student’s t-test, HD versus control. In panels A, B and D, the blue dashed line indicates no CC alteration relative to controls. The complete list of TF families and their CC analyses is provided in Supplemental Table 1.

Next, to determine whether HOXs gain dominance in vulnerable neurons beyond the striatum, we constructed and analyzed the networks of pyramidal neurons from the motor cortex of HD patients and control subjects using the GSE233387 transcriptomic dataset.^5^ A large gain of dominance of the HOXs is observed in vulnerable SATB2-high neurons (Figure 1D), a population characterized by the high expression of the transcription factor SATB2, compared with cells which appear resilient in HD, including SATB2-medium and -negative neurons.^5^

In summary, the gain of dominance of the HOX family is common to vulnerable neurons across brain regions and patients in the datasets we analyzed. In comparison, alteration in the dominance of individual TFs varies significantly across vulnerable neuron types and brain location (Supplemental Table 2).

### HOX dominance correlates with age at disease onset and CAG-Age Product score

After determining HOX dominance is a common denominator across HD brains, we next explored its relationship to disease onset. Ages of motor and cognitive symptom onset and inherited CAG-repeat length were recorded for 21 patients in the nemo:dat-ztfn3cc dataset from whom sufficient cells had been sequenced to build TF networks. We binned data by decades across age ranges from 20 to 80 years and plotted HOX CC in MSNs with age of onset at the time of donation (Figure 2A). This approach provided interpretable age categories while reducing sensitivity to individual variability in the relatively small cohort. Results show HOX CC correlates with the age range of motor and cognitive symptom onsets, with higher CC corresponding to earlier age of onset. The relationship was well described by an exponential fit, with coefficient of determination (R2) of 0.77 and 0.76 for motor and cognitive onsets. While HOX CC is measured in networks from the time of donation, the same relationship is anticipated for the HOX CC at the time of disease onset as HOX CC values correlate directly with disease grades (see below and Figure 3A). For comparison and as a positive control for the analysis, inherited CAG repeat length showed correlations with age of onset of a similar magnitude in the same dataset (Figure 2B). Exponential fits yielded R^2^ values of 0.79 and 0.68 for age at motor and cognitive onset, respectively.

**Fig. 2.**
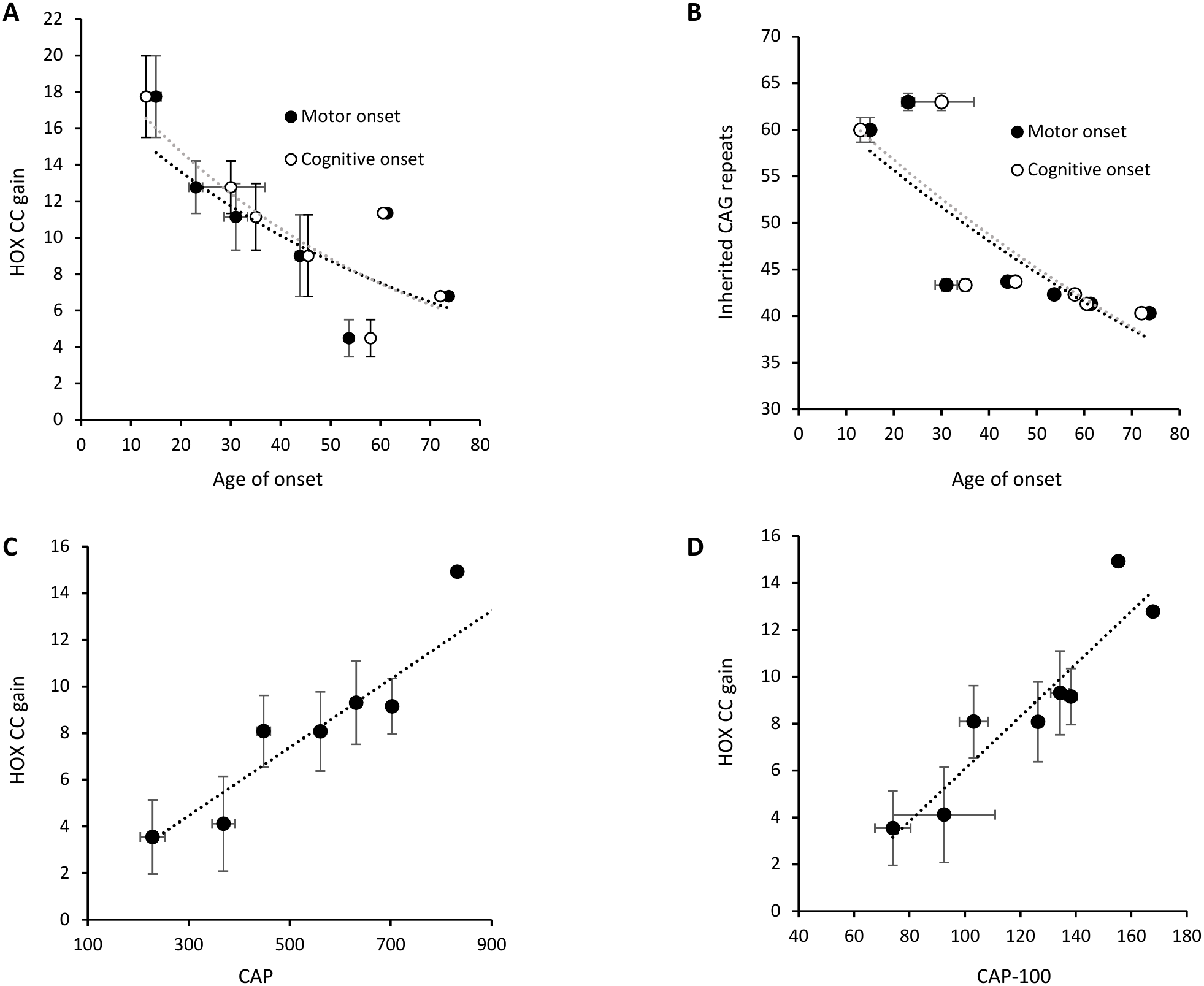
HOX CC gain in HD correlates with age at disease onset and CAG-Age Product (CAP). (A) Mean HOX family CC in MSNs at the time of donation as a function of age at motor and cognitive disease onset. Donors were grouped into decade age-at-onset bins, and mean HOX CC was plotted against the mean age at onset within each bin. Exponential fits yielded coefficients of determination R^2^ of 0.77 and 0.76 for motor and cognitive onset, respectively (N = 21). (B) Corresponding analysis of inherited CAG-repeat length as a function of motor and cognitive age at onset, using the same decade bins (N = 21). (C, D) Relationships between HOX CC at the time of donation and CAP (C) or standardized CAP (CAP-100; D), two measures of cumulative disease burden (N = 38). Data are presented as mean ± SEM, nemo:dat-ztfn3cc dataset.

**Fig. 3.**
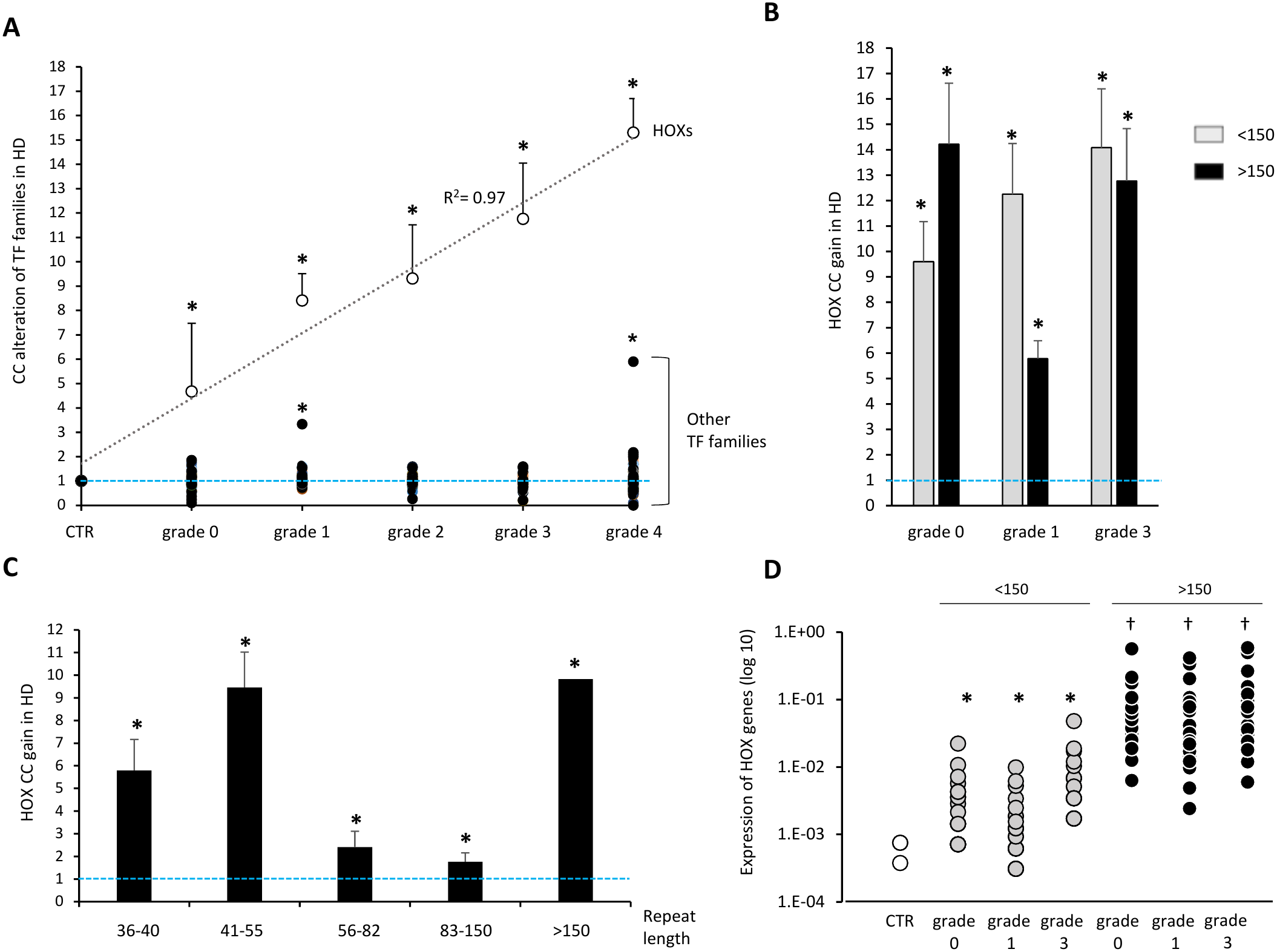
HOXs are functionally dominant across all HD grades and at low and high CAG-repeat lengths. (A) Changes in the CC of human TF families in MSNs across HD grades relative to controls, presented as mean ± SEM. HOX CC gain is shown as white circles and increases linearly with disease grade, R^2^ = 0.97. CC alterations of the NEUROG family are shown as grey circles, and those of the remaining 98 TF families as black circles. For each TF family, CC values in HD were normalized to the corresponding controls within each dataset (GSE152058, grades 2-4; nemo:dat-ztfn3cc, grades 0-4), and the normalized values were combined for analysis. (B) HOX CC gain in MSNs from HD donors with 36-150 or >150 CAG-repeats at disease grades 0, 1, and 3, for which sufficient cells were available for TF-network construction. Data are presented as mean ± SEM. (C) HOX CC gain in HD MSNs across increasing CAG-repeat-length bins, presented as mean ± SEM. (D) Absolute expression levels of HOX TFs in MSNs from controls and HD donors with 36–150 or >150 CAG-repeats. For controls, only HOX TFs with non-zero CC values are shown. Arbitrary units, statistical significance: *p < 0.05 versus controls; †p < 0.05 versus both controls and corresponding CAG <150 groups (Student’s t-test). Datasets used for building networks of Panel A were GSE152058 and the nemo:dat-ztfn3cc dataset, and the nemo:dat-ztfn3cc dataset only for panels B-D. N = 58 controls, 38 HD donors for panel A; for panels B-D, analyses were made for each individual HOX TF present in networks from the 1 control and 8 HD donors with CAG-repeat data from the nemo:dat-ztfn3cc dataset and data were pooled by bin and grade conditions. The blue dashed line indicates no CC alteration relative to controls in panels A-C.

After determining that HOX dominance correlates with age at disease onset, we next examined whether it correlates with the CAG-Age Product (CAP) and standardized CAP (CAP-100) measures of disease burden and HD progression. We plotted HOX CC in MSNs at the time of donation against CAP and normalized CAP-100 scores for the 38 HD donors in the same dataset from whom TF networks could be constructed. HOX CC showed a positive linear relationship with both CAP and CAP-100 (R2 ≈ 0.86 for each; Figure 2C, D); exponential fitting did not improve the correlation.

### HOX dominance is present at grade 0 and increases through grade 4

To further characterize the relationship of HOX dominance to disease progression, we next examined how HOX CC values evolve across disease grades. HD progresses through five grades numbered 0-4 following the Vonsattel classification.^30^ Grade 0 brains appear macroscopically normal but may show 30–40% loss of caudate neurons without prominent astrogliosis; grade 1 is associated with approximately 50% neuronal loss, whereas grade 4 is characterized by severe striatal atrophy, gliosis, and approximately 95% loss of neostriatal neurons.^30,31^ TF networks were constructed from control donors and donors with HD Grade 0 to 4 with sufficient cells sequenced from both the GSE152058 and nemo:dat-ztfn3cc datasets.^28,29^ Combined analysis of HOX CC across 96 donors from the two datasets (58 controls, 38 HD) shows HOXs dominance is already significant at grade 0, and increases linearly through grade 4, the grade with the most pronounced degeneration, with R^2^ value of 0.97 (Figure 3A). Of note, a similar increase is measured in each dataset independently. The grade analysis shows the NeuroG family of TFs can also gain some functional dominance in HD, although only sporadically. Altogether, this data show that HOX dominance is a common denominator in HD, not only across vulnerable cells, but also across disease stages.

### HOX dominance occurs at low and high CAG repeat lengths

Somatic instability varies among individual cells in HD.^29^ As a result, CAG expansion is asynchronous and not all MSNs show the same CAG repeat number within a given patient’s striatum.^29^ We therefore sought to understand how CAG repeat length influences HOX dominance. To achieve this, we analyzed HOX CC in the networks built from the nemo:dat-ztfn3cc dataset, which contains sequencing and CAG repeat data from individual MSNs from eight HD donors.^29^ The data from those donors include MSNs with measurements below 36 CAG repeats. We excluded these observations and restricted our analysis to cells with measured repeat lengths of ≥36, the lowest associated with the disease.^32^ The measured inherited CAG repeat lengths for the eight donors were: 37, 39, 40, 42 (four donors) and 43.^29^

Prior differential gene expression analysis of the dataset reported that HOX expression levels increase in MSNs once CAG repeat length reaches 150, as a signature of a transcriptomic phase associated with rapid degeneration.^29^ Hence we first examined whether there was a difference in HOX dominance below and above 150 repeats. The number of sequenced MSNs was sufficient to build networks from grade 0, 1, and 3 donors individually using broad CAG-repeat length bins spanning 36-150 repeats, and a separate bin for >150 repeats. Results show HOX dominance is similar in MSNs with CAG-repeat lengths below and above 150 CAG repeats (figure 3B). Network analysis of the two major MSN types, D1 and D2, showed similar HOX dominance, with gains in both of ≈ 11 (D1) and ≈ 9 (D2) over controls.

To further characterize the relationship between HOX dominance and CAG-repeat length, we analyzed cells in smaller, prespecified repeat-length bins. The 36-40 and 41-55 bins captured repeat lengths in the inherited CAG range of the eight donors and were interpreted as representing little or no somatic expansion. The 56-82 bin captured cells with initial expansion beyond the inherited range, while the 83-150 and >150 bins represented ranges previously associated with accelerated somatic expansion and neuronal degeneration, respectively. We combined data across donors to construct networks for each bin because the number of sequenced cells per donor was insufficient for separate analyses.^29^ HOXs dominance was statistically significant in each investigated repeat-length bin, although the magnitude of dominance varied in a bimodal pattern. HOX dominance was greatest in the low (36-40, 41–55) and high repeat ranges (>150) and lower in the middle range (56–82, 83-150) (Figure 3C).

Absolute HOX gene expression counts in the dataset confirm that HOXs are expressed in MSNs with CAG-repeat length between 36 and 150, albeit at a low level, and also confirm their increased expression in MSNs with repeat lengths >150 as observed previously by the authors of the nemo dataset (Figure 3D).^29^ The finding that HOXs are dominant in MSN networks when their expression is low or high suggests that they operate as a multi-member, functional unit that has a low sensitivity to the expression level of its components.

### Ectopic and redundant HOX dominance impairs neurons at early and late stages of disease

The primary function of HOXs in the central nervous system is the specification of the rostro-caudal axis of the embryo during development by driving neuronal identity specification programs and making life-death decisions for cells.^33,34,35^ Subsequently, HOX expression becomes mostly repressed in adult brains.^36,37^ There are 39 human HOX genes spatially organized into four chromosomal clusters A, B, C and D, for which spatial and temporal expression during development follows their positions on chromosomes in a coordinated manner defined as co-linearity.^38^ Hence by definition HOX activity in HD is ectopic.

To probe how HOX family members maintain their dominance across the lifespan of vulnerable adult neurons, we compared the functional dominance of individual HOX members in MSNs across the disease grade and CAG repeat expansion axes (Figure 4). A pattern emerges from the comparisons that supports a coherent, family-type activity: (1) 37 of 39 HOX TFs gain dominance over controls at some point along the two axes, supporting a HOX-subnetwork, collaborative activity, (2) approximately half of the HOXs start to gain dominance at grade 0 and their CC value increases gradually until grade 4, at which point they present high individual CC values, (3) HOX TFs that gain the most dominance over increasing disease grades are the same overall as the HOX TFs that are dominant through CAG-repeat expansion, (4) the HOX TFs dominant at low CAG repeats are also dominant when CAG-repeat length reaches 150. Such a family-based mechanism creates a resilient pathogenic system with built-in redundancy in which the abnormal activity of one HOX can be replaced by another HOX while the family output maintains the same degree of dominance. Lastly, 70% of HOX TFs gaining dominance in MSNs during HD are never expressed in the forebrain during normal development but are expressed in the lumbosacral and thoracic sections of the embryo. Hence HOX toxicity in HD is likely driven by their combined spatial and temporal ectopic dominance in mature vulnerable neurons.

**Fig. 4.**
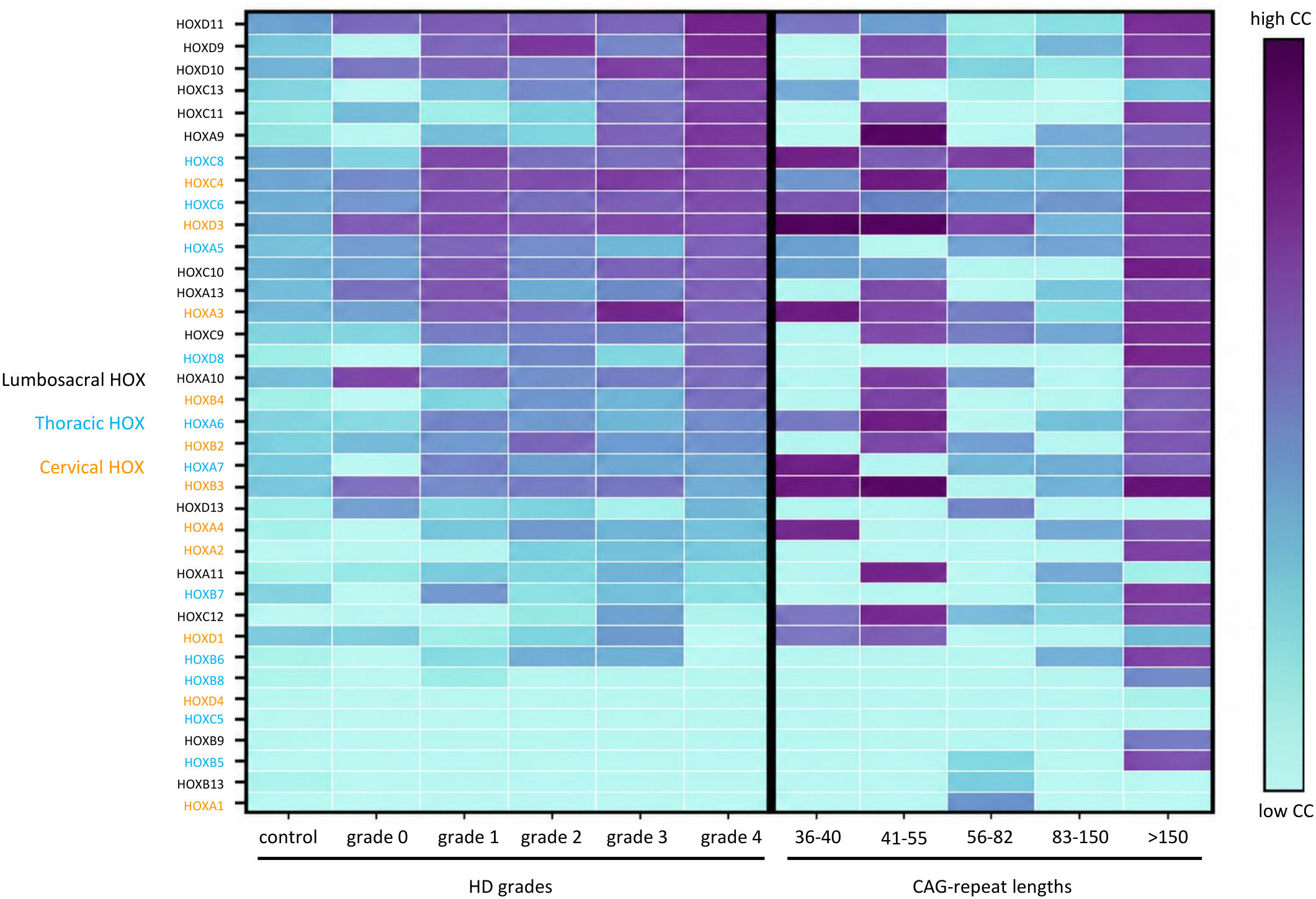
Functional dominance profile of each HOX family member in MSNs as a function of disease grade and CAG-repeat length. The CC value of each of the HOX members present in HD networks is represented in the heatmap as a function of disease grade and CAG-repeat length. The lumbosacral, thoracic or cervical location of each HOX gene expressed during development is color-coded; nemo:dat-ztfn3cc dataset. N = 53 controls and 31 HD donors for which sufficient cells were sequenced to build networks, including 1 control and 8 HD donors with CAG repeat data.

### HOX regulons evolve from dysfunction to degeneration as CAG-repeats expand

Next, we sought to identify the genes directly regulated by HOXs, or HOX regulons, to understand the impact of HOXs on MSN homeostasis. To do so, we constructed entire gene networks of MSNs on top of TF networks from the GSE152058 and nemo:dat-ztfn3cc datasets. Analysis across grades and CAG-repeat lengths shows that over 4,000 genes are regulated by HOXs across two datasets, with 300 genes identified as common HOX targets between the two datasets (Supplemental table 3a). In comparison, HOX regulons in controls only encompass approximately 30 genes. Ontology analysis reveals the universe of HOX-target genes in HD is enriched for intracellular signaling pathways (e.g., calcium, Rap1, Ras, Wnt), neuronal and synaptic identity, cell structure and adhesion, longevity/senescence, cell death and transcription. HOX target genes conserved between the 2 datasets are enriched for calcium and Wnt signaling, cell structure and adhesion, and longevity (Supplemental table 3a).

Comparative ontology analysis of HOX regulons from MSNs with 36–150 versus >150 CAG-repeats provides insights into how HOX-associated transcriptional programs differ across stages of somatic CAG expansion and HD progression at the individual MSN level (Supplemental table 3b). In both groups, HOX regulons retain a broad developmental patterning and neuronal identity signature, encompassing multiple HOX genes, developmental transcription factors including *SIX1, SHOX2, LHX9, ONECUT1* and *TBX5*, and HOX-associated lncRNAs such as *HOTAIR, HOTAIRM1* and *HOTTIP*. Despite this conserved developmental signature, their associated functional networks differ. In MSNs with 36-150 CAG-repeats, HOX regulons show significant enrichment for innate immune and inflammatory pathways, including cytokine signaling, Toll-like receptor signaling and complement activation. These pathways have been implicated in neuroin-flammation and synaptic dysfunction in HD,^28,39,40^ suggesting a potential link between HOX-associated regulatory programs and early neuronal pathology before overt neuronal loss. In MSNs with >150 CAG-repeats, HOX regulons are enriched for neuronal structural and signaling processes, including axon development, synapse organization, Wnt signaling, calcium-mediated signaling, neuronal cell adhesion and postsynaptic cytoskeletal organization. These processes are essential for neuronal identity, connectivity and structural maintenance, consistent with human postmortem transcriptomic studies documenting disruption of MSN identity and neuronal organization during HD progression.^12,29^ Together, these findings suggest that persistent ectopic HOX-associated developmental programs may themselves disrupt mature neuronal identity and homeostasis, while their changing functional associations may introduce additional pathological effects. At lower CAG-repeat lengths, these may involve inflammatory signaling and synaptic dysfunction, whereas at >150 repeats, alterations in neuronal structural organization, connectivity and signaling may further compromise neuronal integrity and survival. This remodeling occurs around the 150-repeat threshold, previously linked to marked transcriptional disruption, loss of neuronal identity and degeneration in HD MSNs, supporting a model in which aberrant HOX-associated regulation may contribute to the progression from neuronal dysfunction to degeneration.

Lastly, the regulon analysis shows that in HD, HOXs form a dense regulatory cluster with a higher-than-normal degree of self-connectivity (>40%). This is highly unusual for a family of TFs, as we measure an average of approximately 1% of intra-member edges for the other major TF families involved in neuron biology, including ETS, KLF, ZBTB or ZSCAN families. Notably, HOX clusters are present at all HD grades and form a near stand-alone HOX-island in the global TF network of MSNs (Supplemental Figure 1). This suggests that HOX family members interact with the broader TF network differently from other TFs and that this pattern is consistent across disease grades.

### HOX dominance becomes independent of mHTT in zQ175 mice

Given that current therapeutic approaches are strongly focused on mHTT-lowering, we next wanted to understand how this therapeutic approach affects HOX dominance. To do this, we applied the network-CC analysis to a dataset generated from zQ175 transgenic mice harboring exon 1 of the human *HTT* gene containing ∼ 175 CAG-repeats (GSE270728). In this study, mice aged 2 or 6 months were treated with vehicle or with the mHTT repressor ZFP30645 which “resulted in robust and selective abrogation of mHTT production >80% in striatal neurons”.^41^ CC analysis of MSN networks of zQ175 animals at different timepoints shows that HOX TFs gain significant dominance in this model compared to wild-type mice (figure 5A). Sustained lowering of mHTT levels with ZFP30645 produced an approximately 55% reduction of HOX dominance in MSNs after 4 months of treatment. However, HOXs regain a similar level of functional dominance over time in MSNs from zQ175 mice treated with the mHTT repressor compared to MSNs from vehicle-treated animals (Figure 5B). This is observed whether the mHTT lowering treatment started at 2 or 6 months of age, showing that HOX network dominance is not fully and durably normalized by substantial mHTT lowering in the model.

**Fig. 5.**
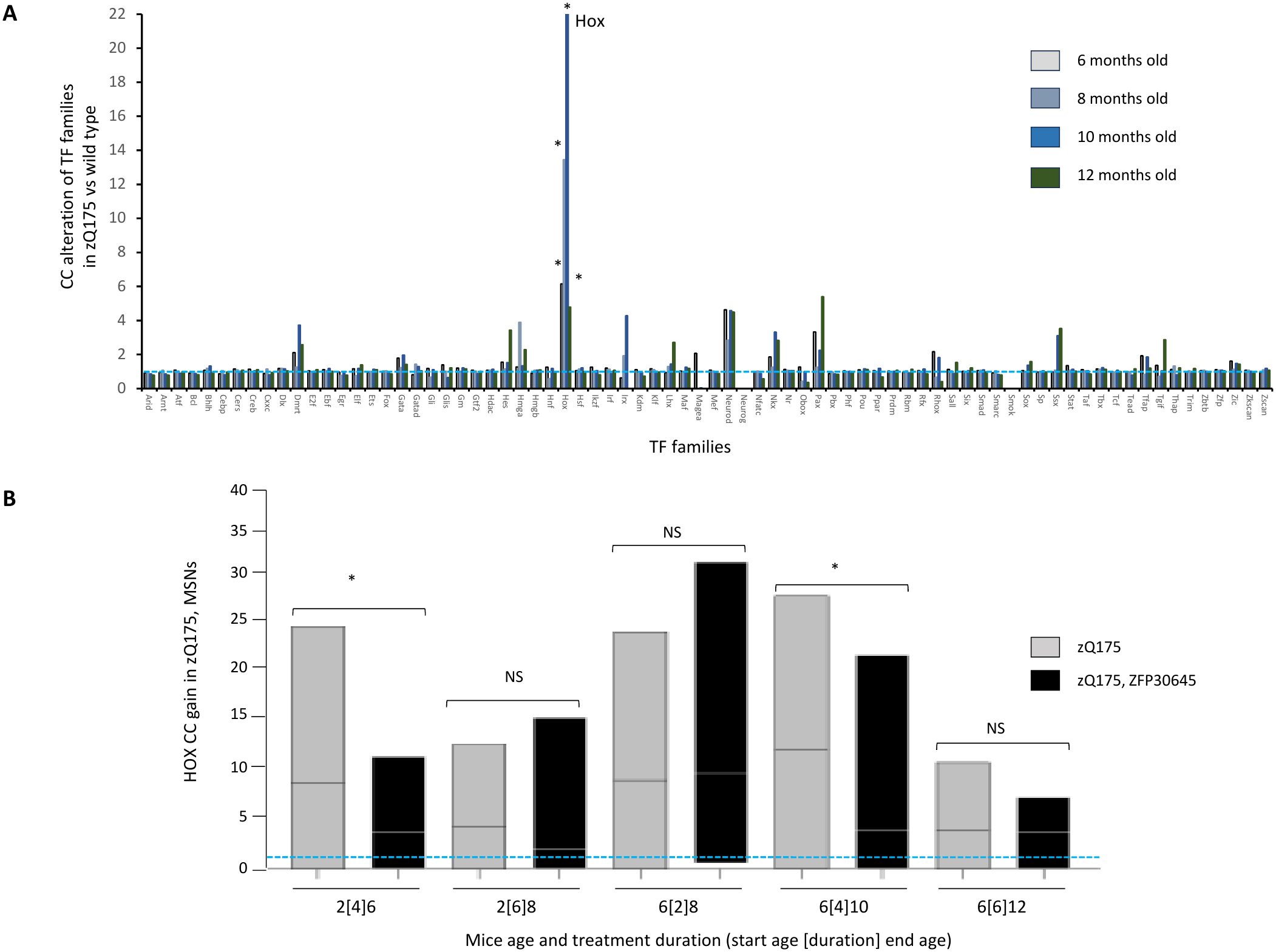
The gain of HOX functional dominance becomes self-sustained in MSNs of zQ175 transgenic mice. (A) Mean CC gain of TF families in MSNs of 6-, 8-, 10- and 12-month-old zQ175 mice; * p<0.05, Student T-test, zQ175 vs wild type. (B) Violin plots of HOX CC with mean (bar), min and max values after 2, 4 and 6 months of mHTT lowering, ZFP30645 treatment starting at 2 and 6 months of age, 2[4]6 = after 4 months of treatment starting at 2 months of age, 2[6]8 = after 6 months of treatment starting at 2 months of age, 6[2]8 = after 2 months of treatment starting at 6 months of age, 6[4]10 = after 4 months of treatment starting at 6 months of age, 6[6]12 = after 6 months of treatment starting at 6 months of age; * p<0.05, Student T-test, zQ175 vs zQ175 ZFP30645, NS = non-significant. The blue dashed line indicates no CC alteration relative to wild type controls. Networks were built from snRNAseq data from the GSE270728 dataset of the Green et al. study.^41^ For analyses, the treatment controls from the study (inactive ZFP-DBD and empty AAV Estop constructs) were combined, as well as MSN groups. Each condition includes at least 9 datapoints.

### COR-H800, an inhibitor of the HOX family

TF network data show that almost all HOX members exert dominance at some point in HD. To inhibit HOX dominance, a drug candidate therefore would need to efficiently block multiple HOX family members simultaneously, and ideally all the HOXs involved at any given disease stage in a given patient. The decoy oligonucleotide technology is well suited to inhibit multiple TFs simultaneously and has shown clinical efficacy in pain and other diseases.^42–47^ A decoy mimics the binding site of its target TF on the genome, and multiple sites can be added to a single molecule.^48^ We designed COR-H800, a decoy that possesses five families of DNA motifs bound by the 39 HOX members. HOXC4, a prototypical HOX that gains functional dominance with each disease grade in persons with HD (Figure 4), was used as a reference to confirm binding activity in a transcription factor ELISA assay (Figure 6A). COR-H800 stability experiments in human cell extracts estimated a high, long-term metabolic stability with no decline measured during the 72h assay (Figure 6B). In mouse brain homogenate, COR-H800 half-life is measured at 15.6 hours (Figure 6B), which reflects the accelerated metabolism that can be observed for oligonucleotides in mice compared to higher species and humans.^49,50^

**Fig. 6.**
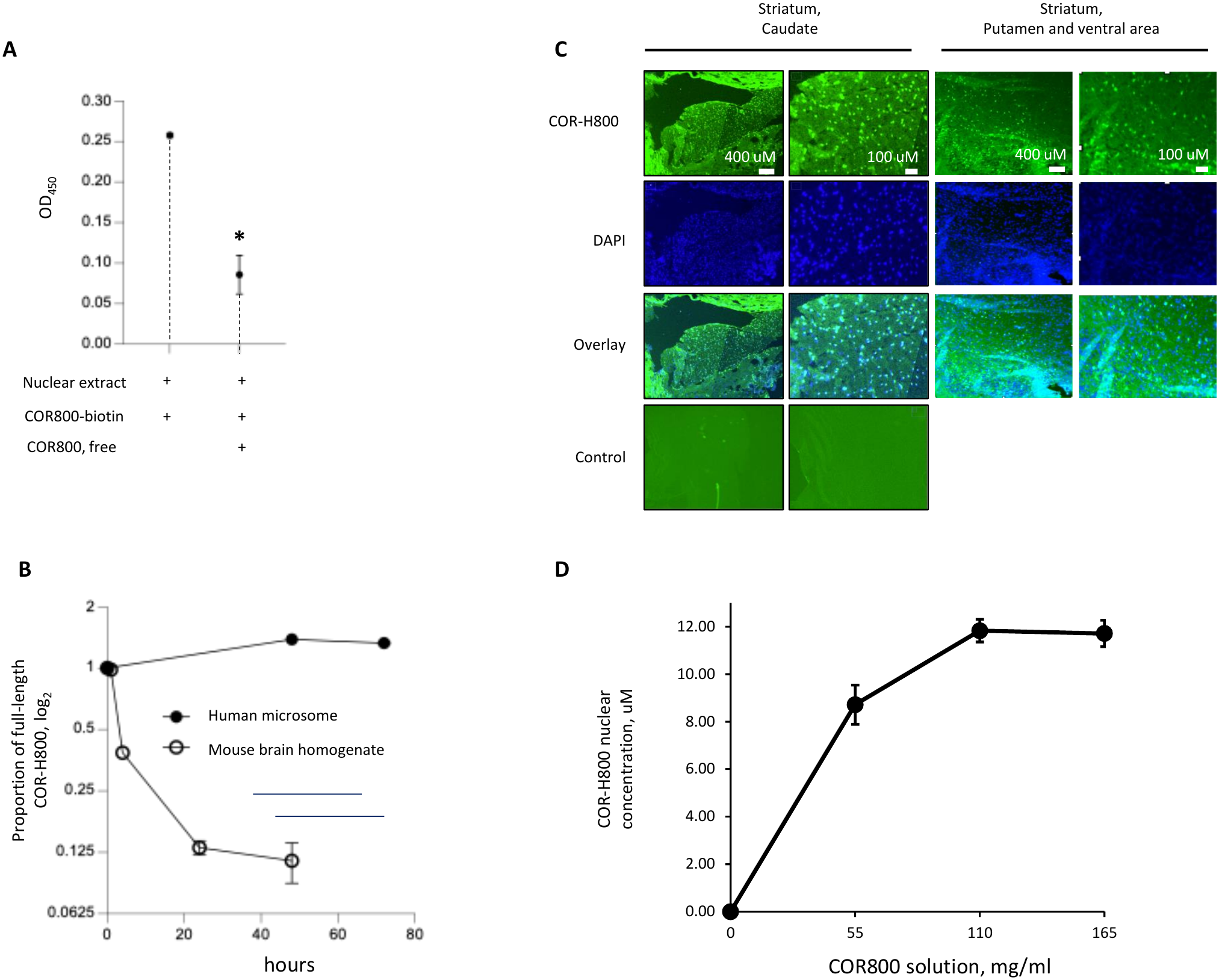
COR-H800 binding, metabolic stability and striatal uptake. (A) COR-H800 HOX binding was confirmed using HOXC4 as a reference HOX member. HOXC4 binding to a biotinylated COR-H800 was measured using a transcription factor ELISA in the presence of 2.5 ug of Jurkat cells nuclear extract and binding specificity was confirmed by adding a competing 8-fold excess of non-biotinylated COR-H800, *p<0.05, Student T-test vs control without free COR-H800, N =3. (B) COR-H800 metabolic stability was measured in vitro by incubating COR-H800 in human liver microsomes and mouse brain homogenates and measuring the proportion of full-length COR-H800 remaining at predefined timepoints, N =3. (C) COR-H800 nuclear localization in the striatum of mice injected with a fluorescein-COR-H800 conjugate and sacrificed 1 hour later. DAPI (nuclear stain) co-staining was used to evaluate nuclear localization. Non-injected mice served as controls for background autofluorescence. (D) COR-H800 nuclear concentration in the caudate region of the striatum as a function of COR-H800 injectate concentration, data presented as mean ± SEM, N = 1-3 mice per condition.

### COR-H800 distributes through the brain following intrathecal administration in mice

Intrathecal decoys distribute deeply into neuronal tissue and naturally enter cells and nuclei.^51^ To confirm COR-H800 brain uptake, fluorescein-conjugated COR-H800 was injected intrathecally at increasing concentration levels in mice. Animals were euthanized 1 h after administration, their forebrains were harvested and processed for imaging quantification. Results show COR-H800 diffuses throughout the forebrain and the striatum at micromolar concentrations (Figure 6C, D, Supplemental Figure 2) when administered at or below 110 mg/ml, a concentration that has previously been effective with intrathecal decoys in the clinic ^52^. Double labeling with DAPI nuclear stain shows that COR-H800 penetrated up to 85% of striatal cells and primarily concentrates in cell nuclei (Figure 6C). COR-H800 nuclear concentration in the striatum increases in a dose-dependent manner and reaches a maximum after injection at 110 mg/ml in the 8-12 micromolar range (Figure 6D).

### COR-H800 inhibits HOX dominance and reprograms TF networks of MSNs in R6/2 mice

COR-H800 was tested in the R6/2 transgenic model of HD with ∼ 160 CAG -repeats to evaluate the effects of blocking HOX activity on HOX dominance and HD-related TF network alterations in vivo. This aggressive R6/2 model progresses rapidly with most molecular and cellular alterations and behavioral symptoms well developed around 8 weeks of age, enabling testing of therapeutic interventions over a period of several weeks.^53^ Four-week-old wildtype and R6/2 mice were treated with vehicle or COR-H800 until they reached 8 weeks of age, assessed for behavioral neurological abnormalities (see next section), euthanized, and their striatum harvested for single-nucleus RNA sequencing and TF network construction.

CC analysis of MSN networks confirmed that HOXs were not functionally relevant in controls and that they gained functional dominance in R6/2 MSNs (Figure 7A). While the HOX gain of dominance measured in HD donors is reproduced in R6/2 mice, it only accounts for ≈ 45% of TF family alterations in these mice (Figure 7A, C) whereas the alteration in HD donors is limited to HOXs. The blockade of HOX activity in R6/2 mice with COR-H800 eliminated HOX dominance (Figure 7B), demonstrating experimentally that HOXs self-maintain their dominance. From a TF network standpoint, a disease arises from profound alterations of a normal network hierarchy in diseased cells.^25^

**Fig. 7.**
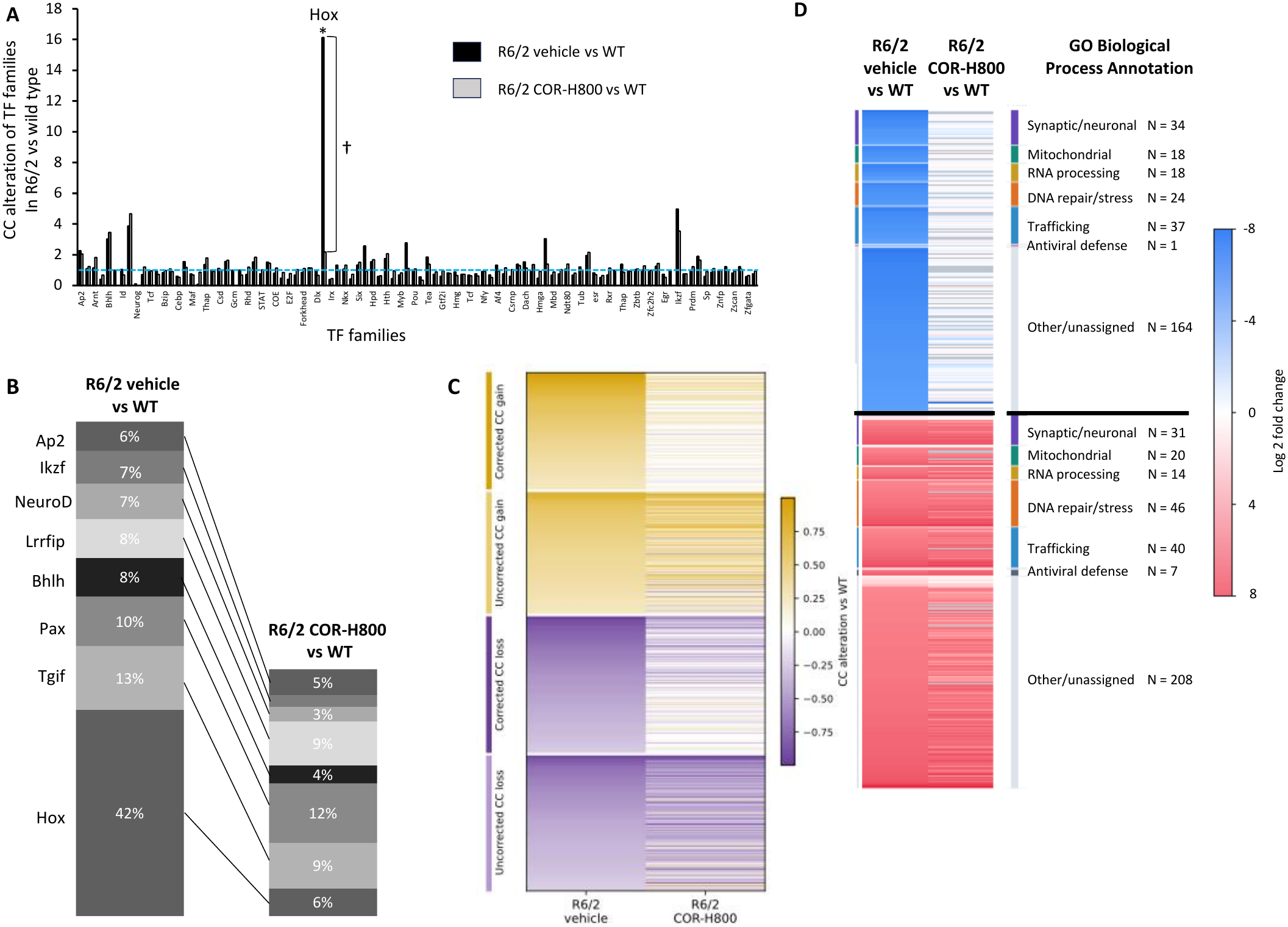
COR-H800 reprograms MSN TF networks in R6/2 mice. (A) CC alteration of TF families in MSNs from R6/2 mice treated for 4 weeks with vehicle or COR-H800 compared to wild-type animals at 8 weeks of age. *R6/2 vehicle vs wild type, †R6/2 vehicle vs R6/2 COR-H800, p < 0.05, Student t-test. (B) Proportion of total CC gain from TF families altered functionally in R6/2 vehicle and R6/2 COR-H800 mice compared to wild-type. For this analysis, families with a CC increase of at least 2-fold in R6/2 vehicle mice vs wild type were considered altered. The total sum of CC gain measured in R6/2 vehicle mice vs wild type for those families was used to normalize the CC gain of each TF family. Results show COR-H800 eliminates HOX dominance in R6/2 mice and restores approximately half of the CC alterations measured for other TF families in R6/2 MSNs vs wild-type. (C) Heatmap showing individual TF CC alterations relative to wild-type in vehicle- and COR-H800-treated R6/2 mice. TFs were classified as exhibiting altered functional dominance when the absolute difference in CC between vehicle-treated R6/2 and wild-type mice exceeded one standard deviation of the absolute CC differences across all TFs. The heatmap shows 667 altered TFs, of which 331 (49.6%) met the predefined correction criteria following COR-H800 treatment. Gold indicates CC gain, purple indicates CC loss, and white indicates CC values approaching wild-type levels. Paired Wilcoxon signed-rank analyses of absolute CC deviations from wild-type showed a 42% average reduction of CC following COR-H800 treatment (p < 0.0001 across all 667 altered TFs). (D) Heatmap of 662 genes differentially expressed in R6/2 vehicle-treated mice relative to wild-type controls, showing changes following COR-H800 treatment. Colors represent log_2_ fold change (blue, downregulated; red, upregulated; white, near wild-type levels; gray, unavailable data). Genes are grouped by selected GO Biological Process annotations. Among the 587 genes with matched measurements, COR-H800 treatment reduced the mean absolute log_2_ fold change relative to wild-type by 39.0%, with reductions of 90.3% for initially downregulated genes and 4.5% for initially upregulated genes. Paired Wilcoxon signed-rank analysis of absolute log_2_ fold changes showed a shift toward WT (p <0.0001, n = 587 genes), WT= Wild-type, N = 1-2 mice per group.

We define network reprogramming as the degree to which the network hierarchy of a diseased cell is restored towards control. Comparison of CC values in MSNs of R6/2 versus wild-type mice shows 51% of TFs have a noticeable gain or loss of CC in R6/2 compared to controls (Supplemental Table 4). Ontology analysis of these TFs revealed significant enrichment for developmental and cell-fate regulatory programs, including neuronal differentiation, cell-fate commitment and pattern specification, together with inflammatory signaling pathways. COR-H800 treatment corrected the CC alterations of approximately 49.6% of these TFs toward wild-type levels (Figure 7C and Supplemental Table 4). These functionally corrected TFs include numerous regulators of developmental patterning, cell-fate determination and neuronal differentiation, indicating that COR-H800 reverses abnormalities in TF functional dominance associated with neuronal identity regulation. Consistent with the HOX-associated developmental patterning signatures identified in human HD MSNs, these findings suggest that aberrant developmental regulatory programs may be therapeutically modifiable, potentially contributing to the restoration of neuronal homeostasis.

To further characterize COR-H800 treatment effects, we performed differential gene expression analysis in R6/2 vehicle- and COR-H800-treated mice relative to wild-type controls. R6/2 vehicle mice exhibited widespread transcriptional dysregulation, with 662 significantly altered genes, including 366 upregulated and 296 downregulated genes (Figure 7D). Ontology enrichment analysis and complementary gene-level GO Biological Process annotation identified concordant functional programs, including RNA metabolism, DNA damage and repair, cellular stress responses, autophagy and intracellular transport (Figure 7D; Supplemental Table 5). Among genes represented in both comparisons, COR-H800 treatment substantially attenuated the disease-associated transcriptional signature, with 97% of genes downregulated and 24% of genes upregulated in R6/2 vehicle-treated mice no longer significantly dysregulated relative to wild-type controls. Annotation of genes showing shifts toward wild-type expression included functions related to synaptic signaling, mitochondrial organization, RNA processing, and protein homeostasis. In parallel, COR-H800 treatment was associated with 197 additional upregulated and 284 additional downregulated genes, alongside two genes reversing from downregulation to upregulation, representing diverse cellular regulatory and metabolic functions. These additional changes may reflect compensatory transcriptional adaptations accompanying the attenuation of disease-associated gene expression, although their functional consequences remain to be established.

### COR-H800 exerted beneficial effects on R6/2 mouse behavior

The wild-type and R6/2 mice underwent behavioral testing at predetermined intervals throughout the 4-week treatment period with COR-H800 or vehicle. The study was powered to detect an effect size of at least 30% in sensitive readouts using four animals per group comprising R6/2 and littermate controls. A detailed testing schedule is presented in Supplemental Table 6. Vehicle-treated R6/2 mice showed low growth rate compared to wild-type animals and progressively developed a cohesive range of neurobehavioral abnormalities matching the model’s reported neurological deficits.^53–55^ Specifically, these animals developed abnormal freezing and vocalization as well as seizures and were impaired in several behavioral assays, including nesting, hindlimb clasping and digging frequency. In contrast, COR-H800-treated R6/2 mice showed a consistent and significant reduction of all these abnormalities (Figure 8, Supplemental figure 3). Wild-type mice remained free of abnormalities throughout the study, regardless of treatment with vehicle or COR-H800.

**Fig. 8.**
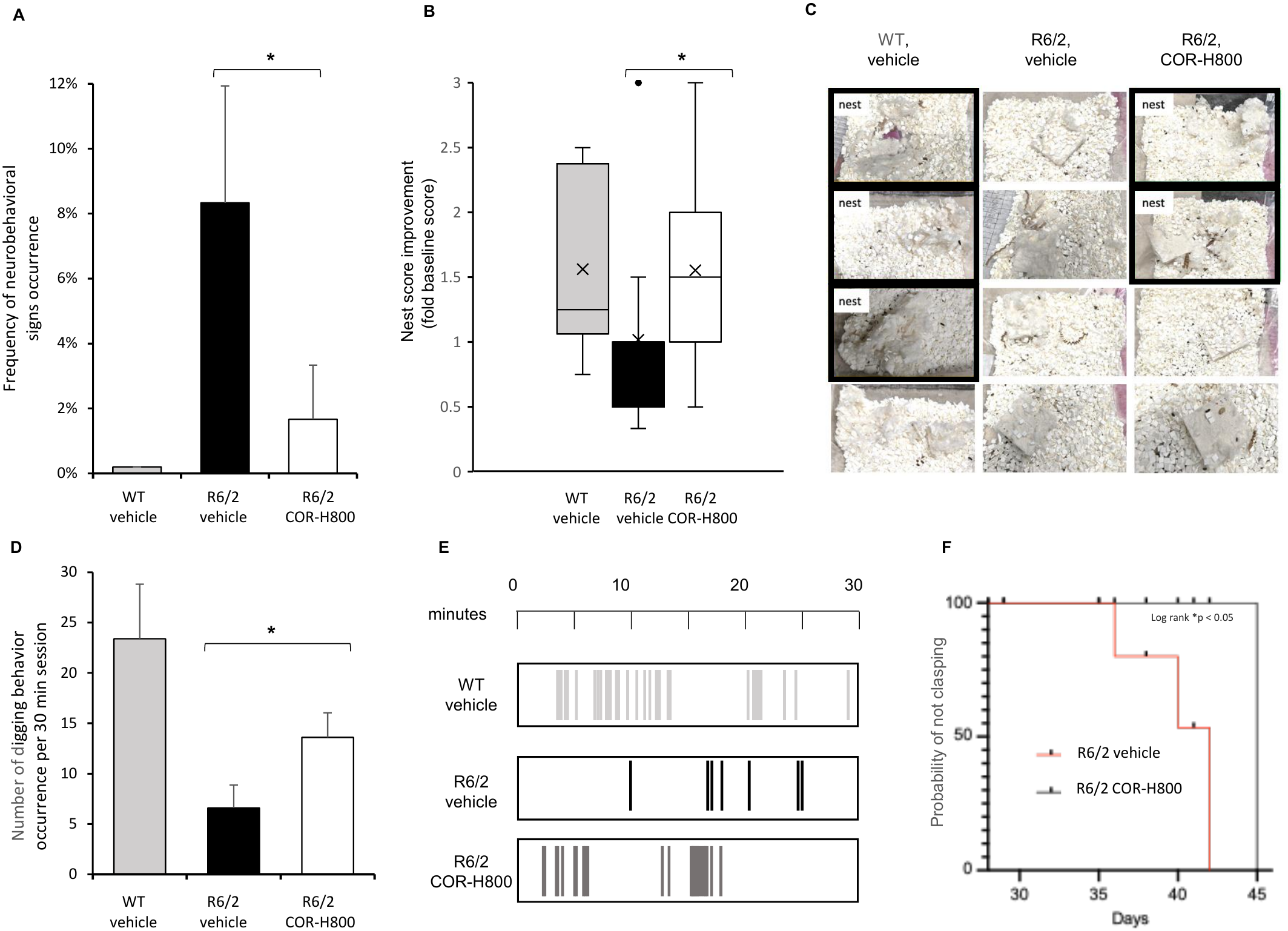
COR-H800 effects on R6/2 transgenic mice behavior. (A) Incidence of neurological signs during the in-cage assessment of the modified Irwin test presented as a percentage of the 15 tests conducted during the study for each group. Signs included abnormal vocalization, abnormal freezing and seizures; * p < 0.05, Student t-test, vehicle vs COR-H800. (B) Nesting capability was measured using a quality nest assessment scale described previously^54^ at baseline and 5 times throughout the study from 4 to 8 weeks of age. Nesting scores were normalized to baseline values and their combined values obtained during the 5 measurements are presented as a box and whisker plot with median (bar) and mean (x). (C) Pictures of nests from each mouse in the presented groups 5 days after vehicle or COR-H800 treatment initiation. Pictures with nests are highlighted in black. No nests were observed from R6/2-vehicle animals. (D) Average occurrence of digging behavior measured in seven 30-minute sessions throughout the 4 week-long study. (E) Digging frequency illustration during a 30-minute testing session. Vertical bars represent occurrences of digging behavior. (F) Kaplan-Meier analysis of time to first clasping showing a longer clasping-free interval with COR-H800 treatment compared to vehicle in R6/2 animals; N = 4 mice per group, WT = Wild type, * p < 0.05, Student’s t-test R6/2-vehicle vs R6/2-COR-H800 for panels A, B and D, *p < 0.05, log-rank (Mantel-Cox) test for graph F.

## Discussion

Transcriptional dysregulation of vulnerable neurons is prominent in HD.^9–12,36^ CC analysis of TF networks shows alterations of individual TF functional dominance in HD, including TFs previously linked to the disease experimentally or through GWAS, such as SREBF2 or ELK factors.^13–19^ However, the functional dominance of individual TFs in HD is variable and depends on neuron type, brain location and disease grade. Beyond individual factors, TFs are known to operate as families to execute or dysregulate cellular functions during development and in mature neurons. A telling example of the latter is the KLF family, where KLF6, 9 and 15 coordinate in the spinal cord to maintain chronic pain.^48^ A TF family mechanism emerges from the analysis of HD networks, in which the ectopic gain of dominance of the HOX family appears as a common denominator to networks of vulnerable neurons. Quiescent in control MSNs, HOXs gain ectopic functional dominance and their CC gain increases linearly with disease grade. HOX CC also correlates with disease onset and CAP markers of disease burden. The emergence of HOX dominance in grade 0 MSNs, before significant overt tissue atrophy, together with increased HOX functional dominance in iPSCs derived from HD donors with 72 and 180 CAG-repeats (HOX CC gain in HD ≈ 6 over control iPSCs, GSE138525), supports the hypothesis that HOX dysregulation is a cell-intrinsic feature of HD pathogenesis rather than a secondary response to tissue atrophy. Together, these findings identify HOX CC as a molecular correlate of HD severity and progression and ectopic HOX activity as a disease mechanism to address therapeutically.

HOX regulons encompass thousands of genes across categories associated with neuronal impairment and degeneration, supporting the toxicity of HOX dominance in vulnerable neurons. The mechanistic importance of HOXs was confirmed experimentally with COR-H800 in the R6/2 transgenic model of HD. HOXs show both similarities and differences between this mouse model and human HD brains: HOXs gain dominance in R6/2 mice relative to controls but account for only approximately half of TF network alterations driven by TF families in mice. COR-H800 treatment was sufficient to restore nearly half of TF network alterations in the MSNs of R6/2 mice toward the wild-type state, a magnitude of effect consistent with the proportion of HOX-controlled network alterations in the model. Importantly, these changes were accompanied by improvements in behavioral deficits. Although the sample size per group was intentionally limited to optimize animal usage and enable testing of wild-type and R6/2 littermates on the same days and thus maximize testing sensitivity, the consistency of COR-H800 effects across all behavioral assays provides experimental support for the benefit of blocking HOX activity in an HD model.

HOX CC increases linearly with disease grade but exhibits a non-monotonic, bimodal relationship with CAG-repeat length, with peaks in the inherited range and above 150 repeats. This suggests that HOX dominance is an early feature of HD that persists across disease stages and is further amplified in degenerating MSNs. Rather than tracking CAG length linearly, HOX dominance appears to reflect an early disease-associated regulatory state that is reconfigured at the high-repeat, degenerative stage. This interpretation is further supported by the comparative analysis of HOX regulons in MSNs with 36-150 versus >150 CAG-repeats. Both regulon populations retain a common developmental regulatory signature, while additional functional programs may modulate their pathological consequences. At lower repeat lengths, regulon genes are predominantly associated with immune and inflammatory signaling, potentially contributing to neuronal dysregulation and impaired homeostasis. At >150 repeats, additional associations with synaptic organization, cell adhesion, and neuronal structural maintenance may compromise neuronal identity and integrity, contributing to degeneration. Together, these findings support a model in which persistent HOX-associated developmental dysregulation, modulated by distinct accompanying programs, contributes to the progression from neuronal dysfunction toward neurodegeneration, consistent with independent evidence that MSN dysfunction precedes neuronal death and with reports linking >150 somatic CAG-repeats to rapid degeneration.^6–8,29^

The difference between the grade-dependent and CAG-dependent patterns may reflect the distinct populations captured by the two analyses. Disease-grade networks integrate MSNs with heterogeneous CAG-repeat lengths, whereas CAG-bin networks are constructed from MSNs with similar repeat lengths. Thus, the progressive increase in HOX dominance across pathological grades may partly reflect changes in the relative representation of these MSN populations, potentially including a declining proportion of intermediate-repeat MSNs and an increasing proportion of MSNs with high CAG-repeat lengths as disease advances.

Overall, CC analysis indicates that HOX dominance is not simply a function of HOX expression level since comparable CC peaks occur at both baseline and elevated expression levels. This is consistent with prior work showing that TF dominance is a function of network hierarchy position rather than transcript abundance.^25^ Dominance does not by itself define a TF’s biological role but rather reflects its functional leverage over one or more cellular functions. The combined CC and regulon analyses therefore converge on a model in which, at similar dominance levels, HOX output shifts from dysregulation at low repeats to degeneration at high repeats.

The bimodal functional pattern suggests two distinct triggers of HOX dominance. The early trigger occurs at low, baseline HOX expression and thus may be post-transcriptional, possibly depending on post-translational modifications or cofactor availability. The late trigger is more likely linked directly to the increased HOX expression observed above 150 repeats, consistent with the reported relief of epigenetic repression by mHTT.^29,56^

Overall, this body of work supports the hypothesis that HOXs are toxic effectors in HD that can uncouple themselves from mHTT and CAG-repeat expansion beyond inherited length. First, HOXs are dominant in the range of inherited CAG-repeat length before significant expansion occurs. Second, analysis of independent zQ175 transgenic mice shows that sustained mHTT lowering only temporarily limits HOX dominance. Although this phenomenon was observed in mice, the extensive self-regulation among HOX family members in human postmortem networks suggests that a similar mechanism may also be present in people with HD. Once induced, HOX activity may be sustained by mutual regulation of HOX-family members. COR-H800 data in R6/2 mice show that the direct inhibition of HOX transcriptional activity is sufficient to eliminate HOX functional dominance in diseased animals, further supporting such a self-regulating mechanism. This HOX behavior is consistent with transcriptional memory, a phenomenon in which trigger-induced transcriptional states persist after the initiating trigger is reduced or withdrawn.^20–24^ Such transcriptional memory has been described in normal and pathological states, including development, pain, inflammation, or autoimmunity.^20–24^ This work suggests HD is also prone to a similar transcriptional memory mechanism and identifies HOXs as contributors to HD pathogenesis and novel targets for disrupting self-reinforcing disease processes.

## Conclusion

This study is, to our knowledge, the first to interrogate HD pathogenesis through functional transcription factor network biology and to identify HOXs as common denominators of the network dysregulation observed in the disease. Ectopic, HDspecific HOX activity in vulnerable neurons appears toxic and data support a significant degree of independence from mHTT abundance and CAG-repeat length expansion beyond the inherited length. Pharmacological blockade of HOX activity with the COR-H800 HOX inhibitor in a transgenic HD model yielded measurable functional benefits in vivo. Altogether, this body of data supports the therapeutic potential of targeting HOXs in HD and warrants further testing.

## Acknowledgements

The authors are grateful to Leora Fox, PhD, of Fox Scientific Consulting, LLC, who provided instrumental editorial support and guidance. This work was funded by Core Biotherapeutics.

## Materials and Methods

### Transcriptomic datasets, network construction and CC analysis

Publicly available transcriptomic datasets were used to construct TF networks: GEO accession numbers GSE152058, GSE233387, GSE270728, GSE180928, GSE138525, GSE138121, GSE108474 and NeMO accession number nemo:dat-ztfn3cc. Methods for constructing TF networks and for conducting CC analyses are described elsewhere.^25^ A minimum of approximately 100-120 sequenced cells from scRNAseq or snRNAseq datasets is required to build networks for a given condition in our hands.

### Test and control articles

COR-H800 decoy (sense, 5′-TTTTACGAGTAATTAGTAAT-TATAATTAATGACCAATAAAA-3′, antisense, 5′-TTTTATTG-GTCATTAATTATAATTACTAATTACTCGTAAA-3′) was manufactured by Integrated DNA Technologies (Coralville, IA, USA) for ELISA and for fluorescein-conjugated COR-H800 brain distribution experiments, and by Synoligo Biotechnologies (Morrisville, NC, USA) for the R6/2 study. For animal dosing, COR-H800 and vehicle were isotonic solutions at neutral pH containing 10 mM Tris, NaCl, CaCl_2_, MgCl_2_, KCl and 0.5% dextrose. Articles were administered intrathecally by percutaneous injection at the cisterna magna level. This site was selected to accommodate repeat dosing in young mice, in which access to the lumbar intrathecal space is restricted. The protocol for intrathecal injections was described elsewhere.^57–59^

### COR-H800 affinity and metabolic stability

Binding of COR-H800 to HOXC4 was assessed using an adapted TransAM Flexi transcription factor ELISA customized for HOXC4 (Active Motif, Carlsbad, CA, USA; catalog no. 41098). A custom biotinylated COR-H800 probe was incubated with Jurkat cell nuclear extracts (Active Motif) in the supplied binding buffer. The probe–protein complexes were subsequently captured on a strep-tavidin-coated microplate, and unbound material was removed by washing. Bound HOXC4 was detected with a rabbit polyclonal anti-HOXC4 antibody (Invitrogen, Thermo Fisher Scientific; catalog no. PA5-100561) at 1:500 dilution. Detection was performed using the kit-supplied horseradish peroxidase-conjugated anti-rabbit secondary antibody and colorimetric developing solution. Following addition of stop solution, absorbance was measured at 450 nm. Binding specificity was assessed by adding an 8-fold molar excess of free, non-biotinylated COR-H800 to the binding reaction. A reduction in HOXC4 binding to the biotinylated probe in the presence of the competitor demonstrates binding specificity. Metabolic stability was measured by incubating a fixed amount of COR-H800 in human liver microsomes (Xenotech, Kansas City, USA, catalog no. H2610) or mouse brain homogenate prepared as previously described^51^ for predetermined durations at 37°C. The amount of remaining parent COR-H800 after each incubation was measured by electrophoresis and normalized to the corresponding control COR-H800 value incubated in parallel in DNase- and RNase-free water.

### Animals

Male C57BL/6 mice, 1-2 months old, were used for the brain distribution study. Male heterozygous R6/2 mice expressing exon 1 of the human mutant huntingtin gene carrying approximately 160 CAG-repeats (B6CBA-Tg(HDexon1)62Gpb/1J strain #:002810) and wild-type littermate controls for the COR-H800 study were obtained from Jackson Laboratory (Bar Harbor, ME, USA). Animals entered the study at 4 weeks of age and were followed for 4 weeks. R6/2 and wild-type mice were drawn from the same litters and were born on the same day. Animal procedures were approved by Stanford IACUC and conducted in accordance with IACUC guidelines and policies.

### COR-H800 brain distribution and nuclear localization

Brain distribution was assessed using a fluorescein-conjugated COR-H800 preparation administered intrathecally. Digital fluorescence images of forebrain sections were acquired using an Axio Imager.M1 fluorescence microscope (Carl Zeiss, Oberkochen, Germany). Fluorescence intensity across the sections was quantified using MATLAB R2025b (MathWorks, Natick, MA, USA) and ImageJ 1.54 (National Institutes of Health, Bethesda, MD, USA). Nuclear localization was assessed by colocalization with DAPI staining. Non-injected animals were used to control for background autofluorescence. Fluorescence measurements were converted to concentration estimates using a standard curve generated by a calibrated scale of fluorescence intensity made by adding 1 uL drops of increasing fluorescein COR-H800 concentration on control brain slides.

### Behavioral testing

Wild type and R6/2 mice were randomly assigned to two treatment groups: vehicle and COR-H800. Four mice were assigned to each group using a computer-generated random sequence with allocation balanced by body weight. All groups underwent treatment and behavioral assessment on the same study days. COR-H800 was administered once at a concentration of 35 mg/ml and then twice weekly at a concentration of 28 mg/ml. Animals were housed at 21-24 °C and 45-55% relative humidity, under a 12-h light/12-h dark cycle, with food and water provided ad libitum. Behavioral assessments were performed blinded. Neurological assessments were performed before and after COR-H800 administration using selected observational components of a modified Irwin assessment.^60^ Mice were observed in their home cages for 15 minutes for abnormalities in spontaneous ambulation, including reduced movement, freezing, or an unsteady gait, abnormal vocalization, and seizures. These signs were also monitored during tail suspension and for 5 minutes after the mice were returned to their home cages. Nesting behavior was measured before treatment and throughout the study using a previously described nest-quality scoring system, with animals housed individually: 1, more than 90% of the Nestlet remains intact; 2, 50–90% remaining intact; 3, 50–90% has been shredded; the cotton is spread around the cage; 4, more than 90% of the Nestlet is torn but the nest is flat; 5, more than 90% of the Nestlet is torn and the nest is a crater.^54,61^ Clasping occurrence and latency as well as digging behavior were measured as described before.^54,55^ A detailed testing schedule is provided in Supplemental Table 6. At the end of the study, animals were euthanized by carbon dioxide inhalation.

### Tissue collection

For fluorescence imaging, brain tissue was immersion-fixed in 4% paraformaldehyde (PFA) in phosphate-buffered saline (PBS) for 72 hours at 4 °C, followed by sequential cryoprotection in 10%, 20%, and 30% sucrose solutions in PBS. Tissue was embedded in Tissue-Tek O.C.T. compound (Sakura Finetek USA, Torrance, CA) at -20 °C and sectioned at 15 µm using a Leica CM1850 cryostat. Sections were collected on slides and coverslipped using VECTASHIELD mounting medium containing DAPI (Vector Laboratories, Newark, CA). Striatal tissue from one wild-type vehicle animal, one R6/2 vehicle animal, and two R6/2 COR-H800 animals was collected for single-nucleus RNA sequencing at the end of the COR-H800 study after 4 weeks of treatment as described previously.^57^

### Single nucleus RNA sequencing

Nuclei were isolated from striatal tissue using the Chromium Nuclei Isolation Kit according to the manufacturer’s protocol (10x Genomics, CG000505). Nuclei concentration and viability were assessed using a Countess 3FL automated cell counter (Thermo Fisher Scientific), with brightfield imaging of trypan blue staining and fluorescence imaging of 7-aminoactinomycin D (7-AAD) staining. Images were analyzed using the instrument’s onboard software. Single-nucleus gene-expression libraries were prepared using GEM-X Universal 3′ Gene Expression v4 chemistry with on-chip multiplexing (10x Genomics, CG000768). Nuclei suspensions were loaded onto a GEM-X OCM chip with reverse-transcription reagents, barcoded gel beads, and partitioning oil. Samples were processed in multiplexing sets of four, with distinct gel-bead barcode sets preserving sample identity. Individual nuclei were partitioned into gel beads-in-emulsion (GEMs), in which polyadenylated RNA was reverse-transcribed using primers containing a 16-nucleotide 10x barcode, a 12-nucleotide unique molecular identifier (UMI), and a poly(dT) sequence. Following reverse transcription, emulsions were broken, and barcoded cDNA was recovered, purified, and amplified. Gene-expression libraries were generated by enzymatic fragmentation, end repair, A-tailing, adaptor ligation, and sample-index PCR, with cleanup and size selection according to the manufacturer’s instructions. Reads from demultiplexed FASTQ files were processed using Cell Ranger multi v10.0.0 (10x Genomics) with the GRCm39-2024-A mouse reference, including intronic reads in gene-expression quantification. Four samples were demultiplexed using their assigned on-chip multiplexing barcodes (OB1–OB4). Across the multiplexed library, Cell Ranger identified 4,768 nuclei, with a mean sequencing depth of approximately 36,688 reads per nucleus.

### Gene Ontology and pathway enrichment analysis

Functional enrichment analyses were performed using ShinyGO (v0.86.0) ^62^ to characterize the biological processes and signaling pathways associated with transcription factors exhibiting altered or COR-H800-corrected CC, as well as genes identified in HOX regulons and differential gene expression analyses. Gene lists were analyzed against Gene Ontology (GO) Biological Process and KEGG pathway databases. Enrichment was assessed using hypergeometric tests with Benjamini–Hochberg correction. Statistical significance was defined as a false discovery rate (FDR) < 0.05. In parallel, individual differentially expressed genes were annotated using mouse GO Biological Process gene sets from MSigDB.^63^ Each gene was assigned a single primary category based on its GO annotations, with genes not matching the selected functional categories classified as other/unassigned. For the comparative analysis of HOX regulons, enrichment was assessed separately for MSNs with 36–150 and >150 CAG repeats.

### R6/2 differential gene expression

Differential gene expression analysis was performed using Seurat V5 in R 4.5.3 within the cell population annotated as SPN neurons.^64,65^ Expression data were normalized using SCTransform, and the SCT assay was prepared for differential expression testing using PrepSCTFindMarkers. Pairwise comparisons were performed using FindMarkers with the default Wilcoxon rank-sum test. Nuclei from the two COR-H800-treated R6/2 animals were combined and compared with those from the wild-type vehicle-treated animal. A separate comparison was performed between the R6/2 vehicle-treated animal and the same wild-type control. Both increased and decreased expression were assessed. Results included average log_2_ fold changes, the proportion of nuclei expressing each gene in each comparison group, and Bonferroni-adjusted p-values. Differentially expressed genes were defined using adjusted p-value threshold of ≤ 0.05.

### Statistical analyses

Comparisons between experimental groups were conducted using Student’s t-tests, and survival-type outcomes were analyzed using the log-rank (Mantel–Cox) test. Correlations were evaluated using linear or exponential regression, with goodness of fit expressed as the coefficient of determination (R2). Gene Ontology and KEGG pathway enrichment were assessed using hypergeometric tests with Benjamini–Hochberg false discovery rate correction (FDR < 0.05). Differential gene expression was evaluated using the Wilcoxon rank-sum test with Bonferroni correction (adjusted p ≤ 0.05). TF CC alterations and their correction by COR-H800 were classified using predefined thresholds. The magnitude of TF CC restoration and differential gene expression recovery was assessed by comparing absolute deviations from wild-type values between vehicle- and COR-H800-treated R6/2 mice using paired Wilcoxon signed-rank tests at the TF or gene level. Data are presented as mean ± SEM where indicated, and sample sizes are specified in the corresponding figure legends.

