## Supplemental Figures for "Reprogramming transcription factor networks with a HOX inhibitor to address Huntington’s disease biology beyond mutant huntingtin"

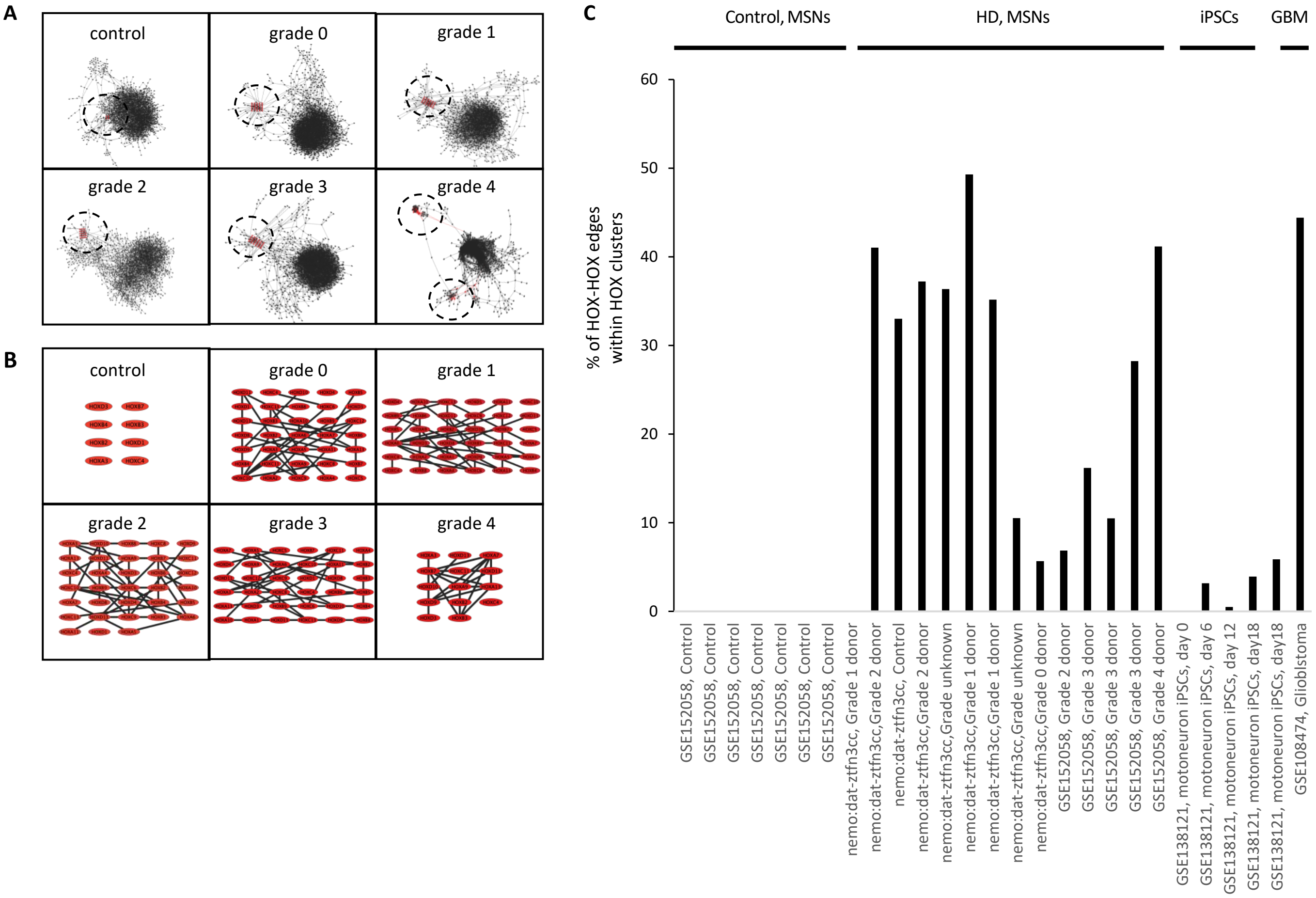


**Supplemental Figure 1. HOX network architecture in HD.** (A) Representative TF networks of MSNs from control and HD donors at each disease grade from the nemo:dat-ztfn3cc (control, grade 0-3) and from GSE152058 (grade 4) datasets. In each network, HOX TFs cluster (circled, red nodes), suggesting a similar way to interact with other TFs at each grade and across patients. Each line or edge represents one TF regulating another TF and each node represents one individual TF. (B) Detailed representation of HOX clusters from (A). Each edge represents an individual HOX member regulating another HOX member. (C) Quantification of HOX clustering measured as the percentage of edges made between HOX TFs vs the percentage of connections made by HOXs with other TFs. Analysis was run in networks from MSNs of control and HD donors from the nemo:dat-ztfn3cc and from GSE152058 datasets, human iPSC motoneurons during differentiation as a model for neuron developmental behavior and in glioblastoma cells (GBM).

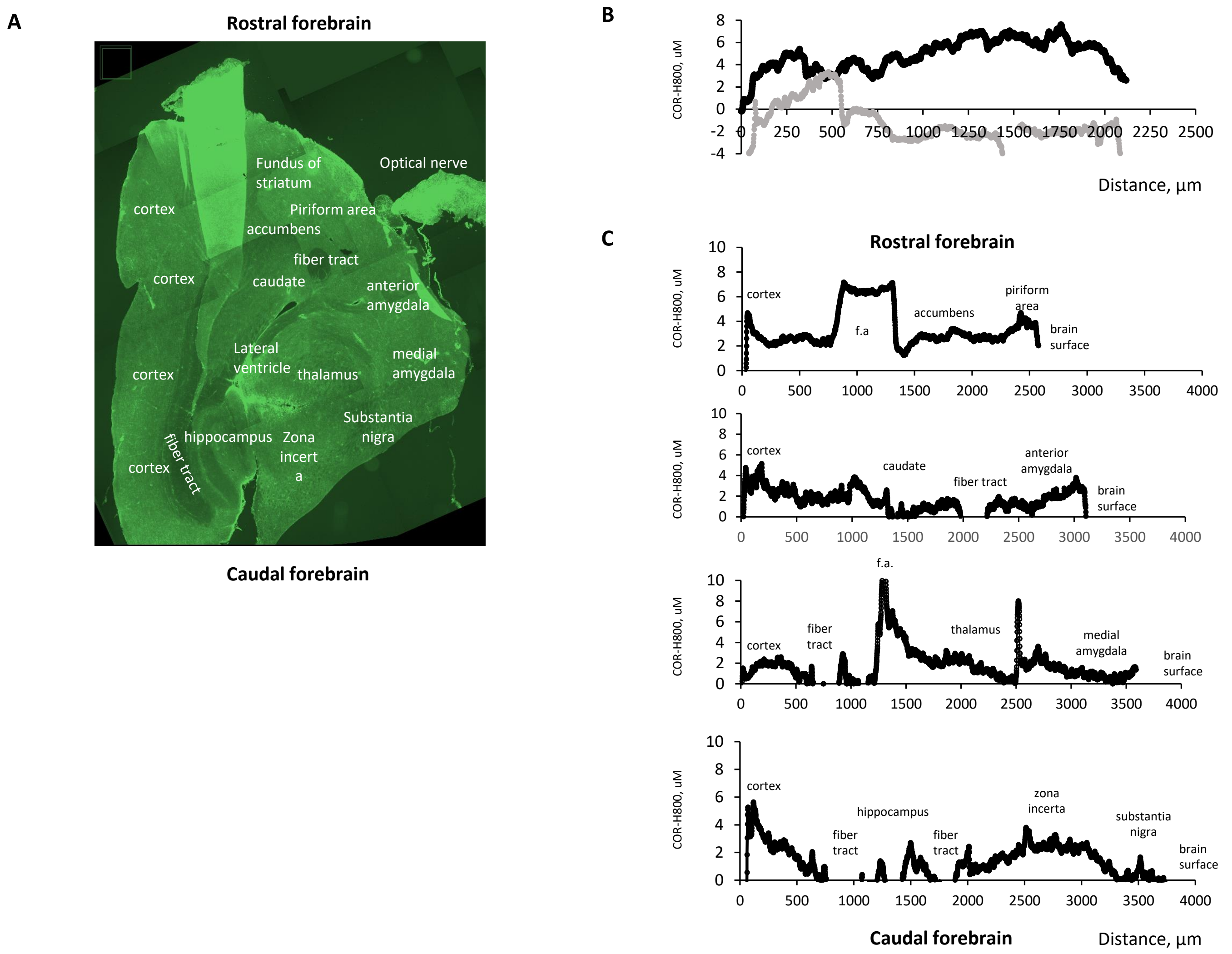


**Supplemental Figure 2. COR-H800 brain distribution.** (A) Mouse forebrain imaging. (B) COR-H800 average tissue concentration (black) across sections vs autofluorescence control (grey). (C) COR-H800 concentration per section from the rostral to the caudal parts of the forebrain of mice. Tissue was harvested 1 h after intrathecal administration of COR-H800 at 110 mg/ml. Distances are measured from the cortex (0 µm) toward the interior part of the forebrain. f.a. = tissue folding artifact.

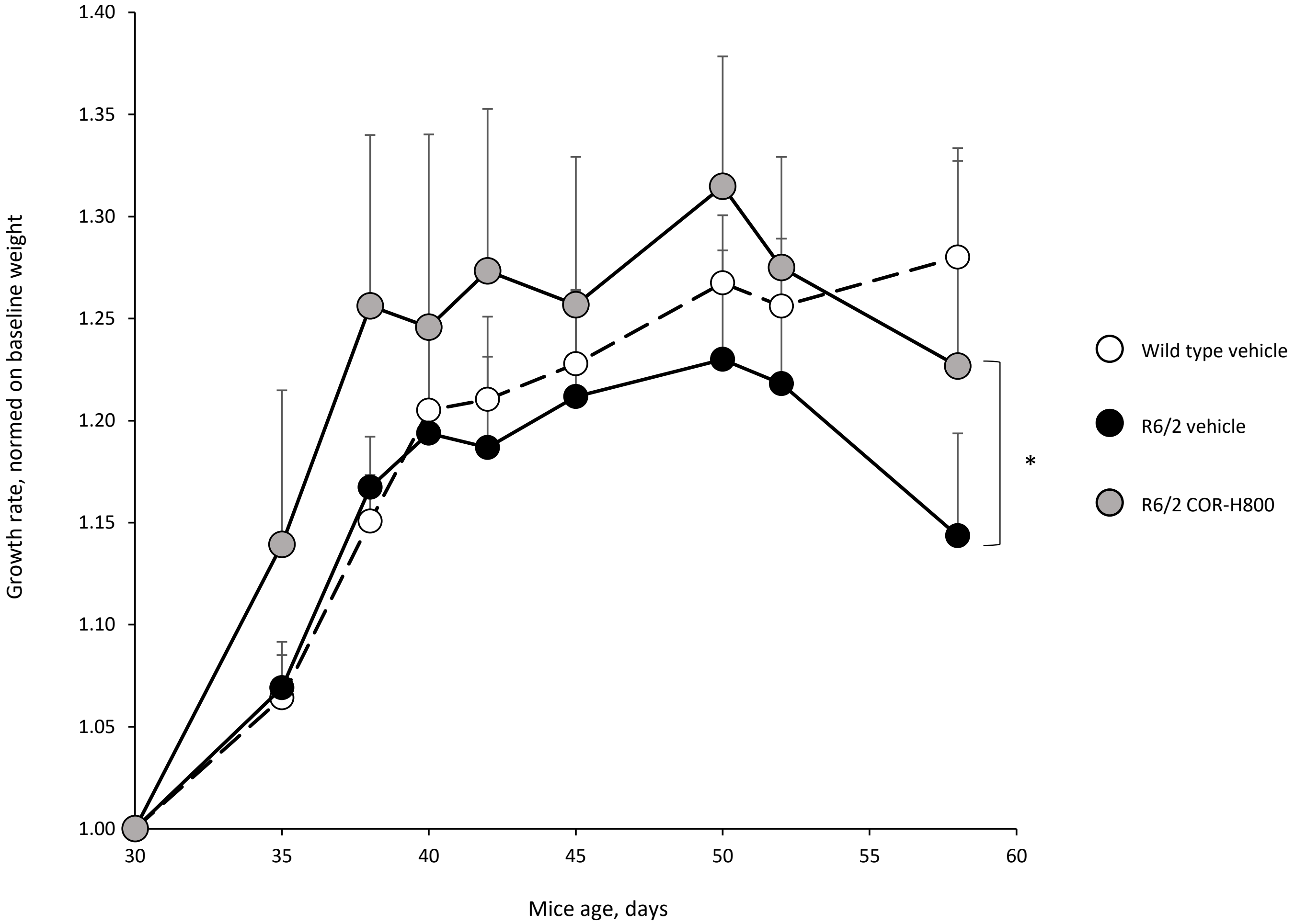


**Supplemental Figure 3. COR-H800 improves R6/2 growth rate.** Animals were weighed at baseline and regularly during the 4-week testing period. Growth rate for each animal was calculated by normalizing its weight at a given time by its baseline value. Data distribution over the testing period, different from vehicle: *p = 0.01, R6/2 vehicle vs R6/2 COR-H800, Student's t-test. Other comparisons were non-significant.
